# The diagonal band of broca regulates olfactory-mediated behaviors by modulating odor-evoked responses within the olfactory bulb

**DOI:** 10.1101/2020.11.07.372649

**Authors:** Inna Schwarz, Monika Müller, Irina Pavlova, Jens Schweihoff, Fabrizio Musacchio, Manuel Mittag, Martin Fuhrmann, Martin K. Schwarz

## Abstract

Sensory perception is modulated in a top-down fashion by higher brain regions to regulate the strength of its own input resulting in the adaptation of behavioral responses. In olfactory perception, the horizontal diagonal band of broca (HDB), embedded in the basal forebrain modulates olfactory information processing by recruiting olfactory bulb (OB) interneurons to shape excitatory OB output. Currently, little is known about how specific HDB to OB top-down signaling affects complex olfactory-mediated behaviors. Here we show that the olfactory bulb is strongly and differentially innervated by HDB projections. HDB-silencing via tetanus toxin light chain led to reduced odor-evoked Ca^2+^-responses in glomeruli of the OB, underscoring HDB’s role in odor response modulation. Furthermore, selective, light-mediated silencing of HDB to OB afferents completely prevented olfactory-mediated habituation and discrimination behaviors. Notably, also social habituation and discrimination behaviors were affected. Mono-transsynaptic tracing studies indicated a robust HDB innervation from the paraventricular nucleus (PVN). Here we provide evidence for a novel tri-synaptic PVN-HDB-OB axis responsible for modulating these types of behavior. Thus, HDB to OB projections constitute a central top-down pathway for olfactory-mediated habituation and discrimination behaviors.

## Introduction

The basal forebrain is considered to play an important role in top-down processing of sensory information (Devore et al., 2016; Lin and Nicolelis, 2008; Sarter and Parikh, 2005; Wilson and Rolls, 1990; Zaborszky et al., 1999). It includes the medial septum, vertical- and horizontal limb of the diagonal band (HDB), the substantia innominate, as well as different peripallidal regions (Zaborszky et al., 1999). The HDB provides significant input into the olfactory network, especially the OB and the piriform cortex (PC) (Gaykema et al., 1990; Gracia-Llanes et al., 2010; Matsutani and Yamamoto, 2008; Niedworok et al., 2012; Zaborszky et al., 1986). Accordingly, classical lesion studies, as well as silencing of distinct neuronal population within the HDB suggested a functional role of this brain region in different olfactory-mediated behaviors (Linster and Cleland, 2002; Linster et al., 2001; Nunez-Parra et al., 2013; Paolini and McKenzie, 1993, 1996; Smith et al., 2015). In this respect, the contribution of the HDB for non-associative memory has been assessed in the context of a simple olfactory habituation and discrimination (dishabituation) paradigm used to probe the neuronal substrate underlying olfactory perception and memory at the initial sensory processing center (Freedman et al., 2013).

Mechanistically, optogenetic stimulation of the HDB resulted in a mixture of excitatory and inhibitory responses within OB (Bohm et al., 2020; Ma et al., 2012; Rothermel et al., 2014; Smith et al., 2015). Importantly, recording of HDB neurons’ response dynamics during either spontaneous odor investigation or odor-reward association learning demonstrated task-related activity modulations (Devore et al., 2016). Collectively, these studies clearly indicated a functional role of the HDB/OB network in olfactory information processing and olfactory-mediated behaviors.

The studies investigating the role of HDB in olfactory-mediated behaviors are complicated by the fact that the HDB significantly innervates, besides the OB, other brain regions involved in olfactory-mediated processing. These regions include the accessory olfactory nucleus, the olfactory tubercle, the piriform cortex, the entorhinal cortex, the orbital cortex, the cingulate cortices, the hippocampal formation and the amygdala (Bloem et al., 2014; Gaykema et al., 1990; Nagai et al., 1982). Thus, silencing the entire HDB, or selected neuronal ensembles within the HDB does not allow to study the functional impact of HDB to OB projections for top-down control of olfactory-mediated behaviors in isolation. To specifically investigate the function of OB-innervating HDB projections in the context of olfactory-driven behaviors isolated from olfactory network effects, we inhibited neuro-transmitter release utilizing inhibition of synaptic release using chromophore-assisted light inactivation (InSynC) (Lin et al., 2013). miniSOG-mediated silencing of presynaptic transmitter release from HDB fibers within HDB resulted in a complete inability of mice to habituate to-, and discriminate between odors. Most notably, we also observed significant deficits in evolutionary crucial social behaviors, including mounting behavior. To identify the underlying neuronal circuits, we performed modified rabies virus-mediated (mRABV) retrograde tracings from HDB and found the hypothalamic PVN to monosynaptically innervate HDB. The PVN, which releases the neuropeptide oxytocin, is known to be critically involved in the regulation of different types of social behaviors. In summary, we revealed a novel PVN-HDB-OB feedback axis involved in the top-down regulation of olfactory-driven behaviors including social behaviors. Most notably, the behavioral effects can be exclusively attributed to an HDB-mediated top-town modulation within OB circuits, emphasizing the relevant role of higher brain regions to directly regulate sensory input strength and thus the adaptation of behavioral responses to olfactory input.

## Results

### Tetanus toxin-mediated silencing of HDB prevents odor habituation and discrimination and decreases glomerular Ca^2+^ activity

It is well established that the OB receives significant top-down input from the HDB (Gaykema et al., 1990; Gracia-Llanes et al., 2010; Matsutani and Yamamoto, 2008; Niedworok et al., 2012; Zaborszky et al., 1986). To further characterize the topology of these projections, we first combined rAAV-mediated neuron labeling and fluoClearBABB-based tissue clearing with 3D-optimized confocal laser scanning microscopy to analyze their distribution within the OB (Schwarz et al., 2015) (Fig. 1A). Stereotaxic delivery of rAAV-Syn-EGFP selectively into HDB and subsequent high resolution 3D-confocal microscopy resulted in an overview of HDB fiber distribution within the OB (Fig. 1B, Suppl. Movie 1). Optical sectioning of this 3D representation demonstrated the differential distribution of the HDB afferents within the different layers of the OB (Fig. 1C). Fluorescent intensity measurements revealed that the GL and GCL are densely innervated by HDB projections, whereas the EPL receives less innervation (Fig. 1D). These results validate the significant monosynaptic innervation of the GL and GCL of the OB by HDB afferents (Niedworok et al., 2012). The robust synaptic innervation of GL and GCL by fibers emanating from HDB emphasize their modulatory potential on olfactory information processing (Aungst et al., 2003).

**Figure 1.**
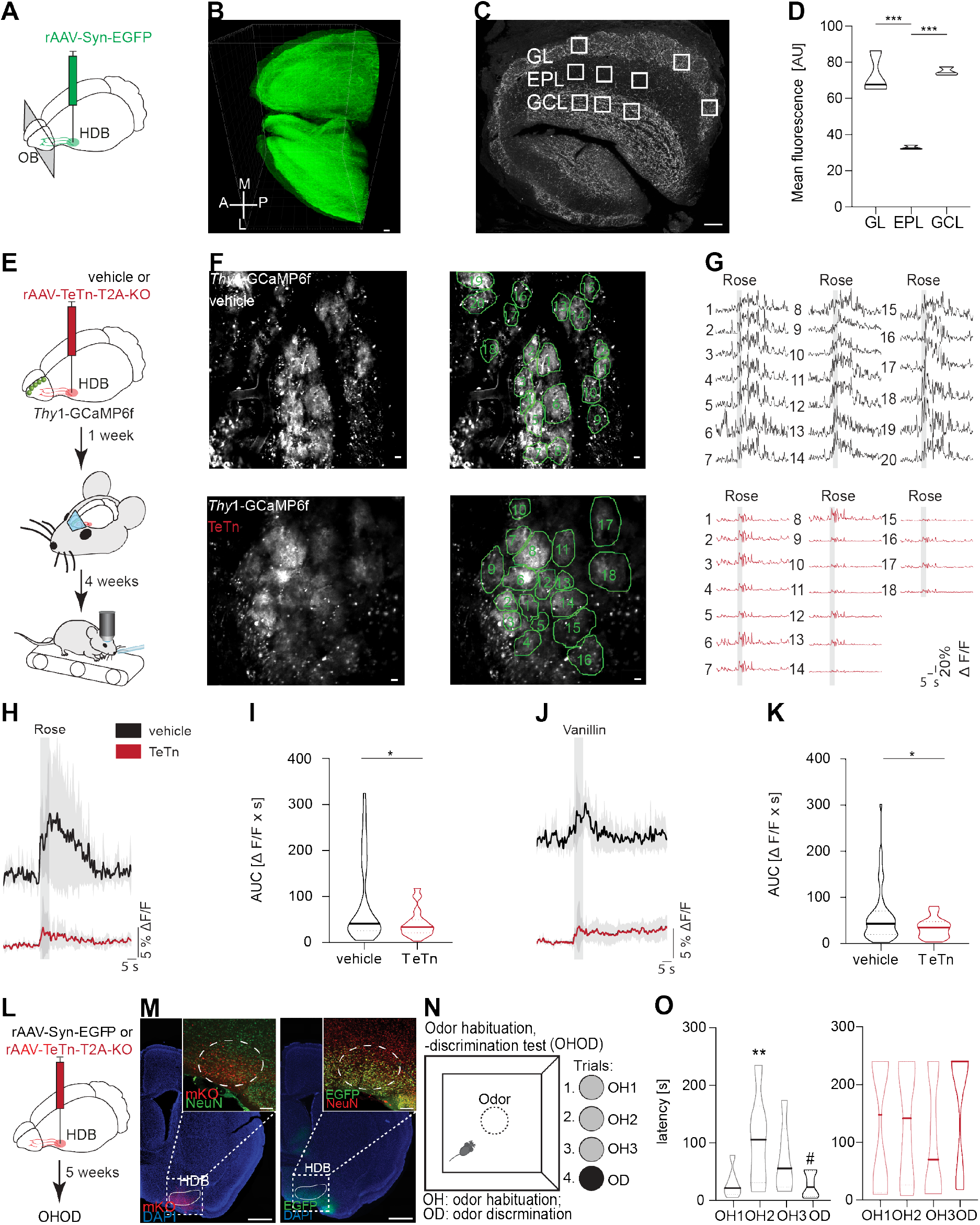
TeTn-mediated HDB silencing decreases GL Ca^2+^-responses and prevents odor habituation/discrimination behavior. **(A)** Schematic of stereotaxic injection. **(B)** 3D representation of afferent HDB fiber occupancy within OB. Scale bar: 100 µm. **(C)** Representative horizontal optical section through the OB showing the different layers used for fluorescence intensity measurements. Scale bar: 100 µm. **(D)** Quantification of fluorescence intensities within the GL, EPL and GCL of the OB. **(E)** Schematic illustration of TeTn-injection into the HDB of *Thy1*-GCaMP6f mice and awake head-fixed Ca^2+^-imaging in the OB during odor stimulation. **(F)** Exemplary average intensity projection of *in vivo* time-lapse of GCaMP6f expressing neurons in the olfactory bulb of vehicle or TeTn-injected transgenic *Thy1*-GCaMP6f mice. Scale bar: 20 µm. **(G)** Ten trial averaged Ca^2+^-transients of ROI1-18 in response to rose odor stimulation in vehicle- or TeTn-injected mice. **(H)** Grand average of all ROIs from n=5 vehicle and n=4 TeTn-injected mice upon rose (2-phenylethanol) odor stimulation. **(I)** Area under the curve of vehicle (n=118 ROIs in 5 mice) compared with TeTn-injected mice (n=107 ROIs in 4 mice). **(J, K)** Analog to (H, I) stimulated with vanillin odor. Mann-Whitney test, *p < 0.0001. **(L)** Schematic of the stereotaxic injection. **(M)** Post mortem analysis of stereotaxic virus injection into HDB. Left: representative coronal section from a TeTn-injected mouse; TeTn infected cells are shown in red (TeTn fused to mKO); close-up in top right corner illustrates the overlap between mKO (red) and neuronal marker NeuN (green). Right: representative coronal section from a vehicle-injected mouse; virus infected cells are shown in green (EGFP); close-up in top right corner illustrates the overlap between EGFP (green) and NeuN (red). **(N)** The odor habituation/discrimination test (OHOD) performed in an open-field arena (60×60cm). The circle in the middle of the arena indicates a petri dish containing a filter paper (3M) with odorant and covered with fresh bedding. Rose and vanillin were used in habituation and discrimination trials. **(O)** Latency (time to reach the petri dish) of vehicle- and TeTn-injected mice during odor habituation and discrimination trials. 1-way ANOVA followed by Dunnett’s posttest (*) and t-test (#) of vehicle-(n=6) and TeTn-injected (n=6) mice. #, *p < 0.05, **p < 0.01. All data presented as mean ±SEM (transients) or median ±quartiles (violin plots). HDB: horizontal limb of the diagonal band of Broca; OB: main olfactory bulb; GL: glomerular layer; EPL: external plexiform layer; GCL: granule cell layer.

To investigate possible alterations in olfactory computations, we silenced the HDB by virally expressed tetanus toxin light chain (TeTn), and subsequently measured glomerular response to odor stimulation using 2-photon Ca^2+^-imaging in awake head-fixed mice (Fig. 1 E, F). For this purpose, we implanted cranial windows above the OB in Thy1-GCaMP6f.GP5.5 mice expressing GCaMP6f in subsets of mitral/tufted cells (M/TCs) (Dana et al., 2014). Olfactory stimulation with two different odors (vanillin and rose) resulted in odor-specific glomerular Ca^2+^-responses in vehicle treated mice (Fig. 1G), whereas in TeTn-treated mice odor-specific responses to both odors were significantly decreased (Fig. 1H-K). Measuring the area under the curve of glomerular Ca^2+^-transients upon rose and vanillin odor stimulation revealed significantly reduced Ca^2+^-responses in mice with TeTn-induced HDB silencing compared to vehicle-injected mice (Fig. 1 I, K). These results suggest that HDB silencing leads to decreased M/TCs response upon odor stimulation in awake mice.

This modulation of M/TC firing after HDB silencing leads to changed OB output, which might alter behavioral responses to odor cues. To probe the effect of HDB on general cognitive abilities as well as odor habituation/discrimination we globally silenced HDB output activity by virally expressed TeTn (Fig. 1, L-M). First, we determined the effect on general cognitive abilities using the puzzle box test (Suppl. Fig. 1K, L). In the puzzle box test mice are required to move from the brightly lit side of the arena to the dark one within a limited amount of time (Galsworthy et al., 2005). The pathway between the two compartments is either an open door with no obstructions, or an underpass, free or blocked. The trials are arranged with increasing complexity (Suppl. Fig. 1K). TeTn- and vehicle-injected control mice showed comparable latencies to enter the goal compartment, with no statistical difference between groups (Suppl. Fig. 1L) suggesting that TeTn-mediated silencing of the HDB does not affect general cognitive abilities tested in the puzzle box.

To further exclude the possibility that TeTn-mediated silencing of the HDB affects odor detection, rendering mice anosmic, we used the buried food test (Alberts and Galef, 1971), in which mice are required to locate a piece of familiar palatable food buried in the bedding of the cage (Suppl. Fig. 1I, J). Both vehicle-control and TeTn groups were able to find and consume the buried food without differences in latency, demonstrating that smelling of an odor in general is preserved in HDB silenced mice (Suppl. 1I, J).

After excluding effects of HDB-silencing on general cognitive abilities and odor sensation, we next investigated its effect on odor habituation/discrimination behavior (Fig. 1N-O). To this end, mice were tested for spontaneous odor discrimination abilities utilizing a simple odor habituation/odor discrimination test (OHOD; Fig. 1N, O; Suppl Fig. 1A-C) (Paolini and McKenzie, 1993). In this task, the same odor is presented to mice over several subsequent trials to assess olfactory habituation. In the last trial, the odor is exchanged with a novel one to test olfactory discrimination (dishabituation). As expected, in the vehicle-control group the latency to approach the odor in the center of the arena increased during the habituation trials and decreased for the discrimination trial. Notably, the TeTn-injected mice did not show significant differences in the latency during odor habituation and discrimination (Fig. 1O). Varied experimental conditions during OHOD lead to comparable results regarding both test parameters, latency and investigation time (Suppl. Fig. 1A-E). Furthermore, previous OHOD test experience including operational procedures did not change the OHOD result in subsequent tests (Suppl. Fig. 1F, G), ruling out a possible effect of prior test experience. Overall, our results demonstrate that TeTn-mediated silencing of the HDB decreases glomerular Ca^2+^-responses and leads to loss of odor habituation/discrimination behavior. However, general cognitive and odor detection abilities remain intact.

### Optogenetic silencing of HDB afferents within the OB prevents odor habituation and discrimination

Apart from the attenuation of the M/TCs responses upon odor stimulation measured in awake mice after global HDB silencing, other HDB afferents may contribute to the severe behavioral deficits during olfactory habituation and discrimination. Thus, in order to selectively study the causal contribution of HDB to OB projections to the observed behavioral deficits in isolation, we inhibited neurotransmitter release from HDB afferents utilizing inhibition of synaptic release with chromophore-assisted light inactivation (InSynC) ((Lin et al., 2013); for details see Methods). To this end, we virally expressed the functional component of InSynC, miniSOG in HDB and chronically implanted light fibers into the OB GCL to trigger chromophore assisted light inactivation (CALI) of presynaptic vesicular release from HDB afferents upon blue light (488nm) illumination (Fig. 2A). Three weeks after virus delivery both vehicle-control-(tdTomato) and miniSOG-expressing mice were illuminated with blue light to silence HDB afferents. To assess if olfactory habituation and discrimination were affected upon synaptic silencing and if this effect could be reversed, we conducted the OHOD test two and 14 days after illumination. Moreover, we repeated the illumination and OHOD with the same group of mice after 8 weeks to test the long-term integrity of miniSOG expressing neurons (Fig. 2A, right side). As shown in Fig. 2B (left panel), neither the light fiber implantation, nor the blue light illumination had effects on olfactory habituation/discrimination indicated by significantly increasing latencies between OH1 and OH3, that had a strong tendency to decrease again in OD in the control group. Importantly, the investigation time for the control group was decreasing between OH1 and OH3 accordingly, and increased again in OD (Suppl. Fig. 2B, left panel). Thus, both parameters, latency and investigation time indicated intact habituation and discrimination abilities in the vehicle-injected control group (note that all three OHOD tests (Fig. 2B, C, D left panels) showed intact odor habituation/discrimination behavior).

**Figure 2.**
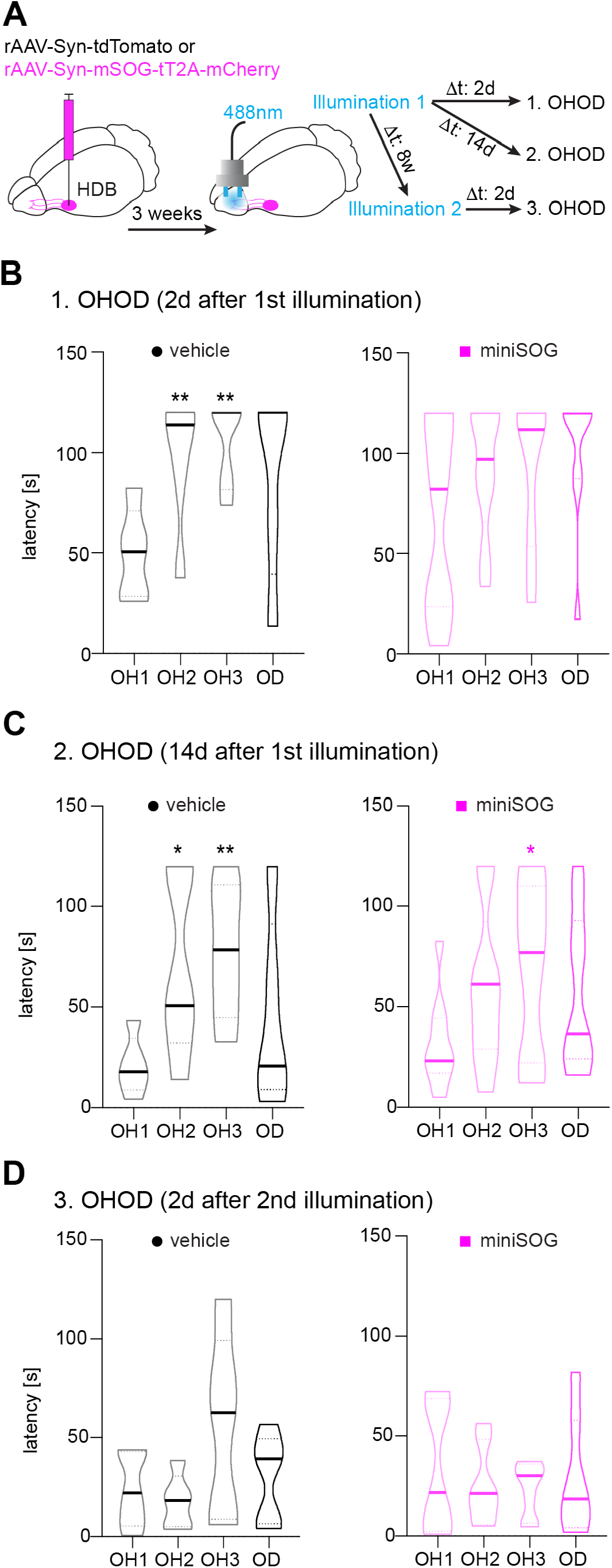
miniSOG-mediated, light-induced silencing of HDB afferents within OB impairs odor habituation/discrimination behavior. **(A)** Timeline of experiment. The odor habituation/discrimination task (OHOD) was performed as already described in Fig 1N, O. **(B-D)** Latency (time to reach the petri dish) of vehicle-injected control and miniSOG-injected mice during odor habituation/discrimination trials 2 days (B) and 14 days (C) after the first illumination, as well as 2 days (D) after the second illumination. Repeated 1-way ANOVA followed by Dunnett’s posttest (OH1 to OH3, (*)) and t-test (OH3 to SD (#)). *p < 0.05, **p < 0.01.

Most importantly, both odor habituation and discrimination were absent two days after blue light illumination in the miniSOG test group, demonstrating that synaptic silencing of HDB afferents within the OB GCL alone is sufficient to prevent odor habituation and discrimination (Fig. 2B and Suppl. Fig. 2B, right panels). Remarkably, the inability to habituate to-, and discriminate between odors was completely restored 14 days after illumination (Fig. 2C and Suppl. Fig. 2C, right panels), but could partially be induced again by repeated illumination after 8 weeks suggested by a tendency for odor habituation/discrimination (Fig. 2D and Suppl. Fig. 2, right panels). Interestingly, we observed a decrease in average latencies during the second test for both groups, suggesting decreased anxiety levels after performing the test for the second time (please compare Fig. 2B with Fig. 2C).

Taken together, our results indicate that complex odor habituation/discrimination behavior depends on the direct modulation of the OB circuits modulated by HDB afferents targeting the OB.

### A novel tri-synaptic basal forebrain circuit linking odor perception with social behaviors

So far, our results clearly indicate an important role of HDB to OB afferents in olfactory-mediated habituation/discrimination behavior. However, since the evidence for a direct synaptic reciprocal coupling between OB and HDB is lacking, it is unclear how olfactory perception is linked to HDB top-down signaling. To identify additional nodes of this pathway, we utilized modified rabies virus (mRABV)-based mono transsynaptic retrograde tracing (Niedworok et al., 2012; Osakada and Callaway, 2013; Wickersham et al., 2007a; Wickersham et al., 2007b; Wickersham et al., 2010) and injected a rAAV/RABV-cocktail (for details, see Methods) into the OB and rAAV-CBA-RG alone into the HDB to allow an additional transsynaptic traversal (Fig. 3A). This approach allowed us to retrogradely trace the neurons synapsing selectively onto the HDB neurons monosynaptically connected to OB interneurons. Surprisingly, this approach revealed spatially confined EGFP expression in the PVN and, to a lesser extent, the supraoptic nucleus of hypothalamus (Fig. 3B).

**Figure 3.**
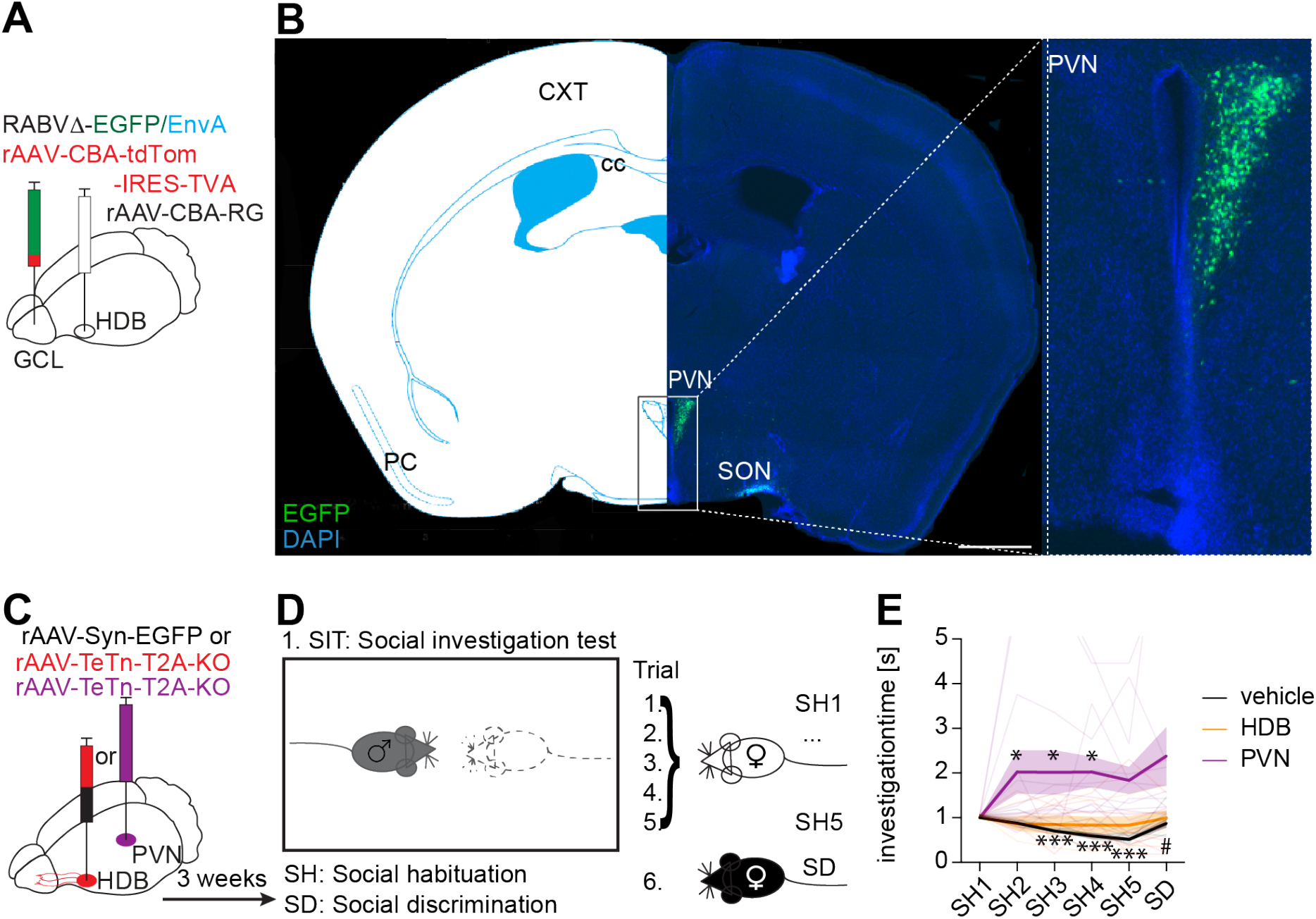
HDB receives direct synaptic innervation from the PVN, and TeTn-silencing of HDB or PVN prevents olfactory-mediated social behaviors. **(A)** Schematic of the stereotaxic injection. Virus cocktail (modified Rabies virus containing EGFP together with supporting rAAVs, see methods) was injected into the OB GCL, while a single rAAV expressing rabies virus glycoprotein (RG) was injected into HDB (n=6 mice). **(B)** Right: Coronal section through the brain at Bregma −0.82mm 10 days after virus delivery. Neurons, back traced from HDB are shown in green. Left: a corresponding scheme obtained from the mouse brain atlas (Franklin and Paxinos, 3^rd^ edition). Inset: Magnification of the region marked in B. **(C)** Schematic of stereotaxic injections into HDB (red) and PVN (purple). **(D)** The social interaction test (SIT) was performed in the male’s home cage. Male mice were presented 5 times with the same ovariectomized female (SH1 – SH5). In the 6^th^ trial a novel female was presented. **(E)** Normalized investigation time of control (black line) and TeTn-injected mice during SIT (yellow/purple lines). Repeated 1-way ANOVA followed by Dunnett’s posttest (SH1 to SH5, (*)) and t-test (SH5 to SD (#)). *p < 0.05, ***p < 0.01.

The PVN is especially known for producing the peptides oxytocin and vasopressin, both important for various aspects of social behavior (Tang et al., 2020). The fact that the PVN is synaptically coupled to HDB suggests a PVN-HDB-OB axis possibly important for olfactory-mediated social behaviors. To functionally probe a causal relationship between the nodes of this newly identified PVN-HDB-OB axis for olfactory-mediated social behaviors, we virally silenced either the HDB, or the PVN with TeTn and tested mice in a social interaction test (SIT) that is in principle analogous to the OHOD paradigm (Ferguson et al., 2000)(Fig. 3C, D). Olfactory-mediated social habituation to a female is reflected by decreasing investigation times upon repetitive presentations (SH1–SH5), whereas social discrimination is reflected by an increased investigation time upon presentation of a novel female (SD, Fig.3 D).

As expected, the vehicle-injected control group showed intact habituation and discrimination indicated by a decrease in investigation time between SH1 and SH5 and an increase during the SD trial (Fig. 3E black line). Most notably, after silencing HDB with TeTn we observed a total loss of olfactory-mediated social habituation and discrimination abilities (Fig. 3E yellow line) (for analogy to HDB silencing in OHOD also see Fig. 1O; Fig. 2 B). Importantly, the PVN-silenced males also did not habituate to the familiar female upon repeated presentations, nor did they significantly increase their investigation time upon presentation of a novel female (Fig. 3E purple line), again indicating a loss of olfactory-mediated social habituation and discrimination.

Importantly, both HDB-silenced, as well as and PVN-silenced males also showed impairments in mounting behavior during the SIT (see Fig. 4A, B). The mounting score reflects the number of trials in which males engaged actively in mounting behavior. Note, that both, HDB and PVN-silenced groups showed significantly fewer mounting attempts (Fig. 4A, B).

**Figure 4.**
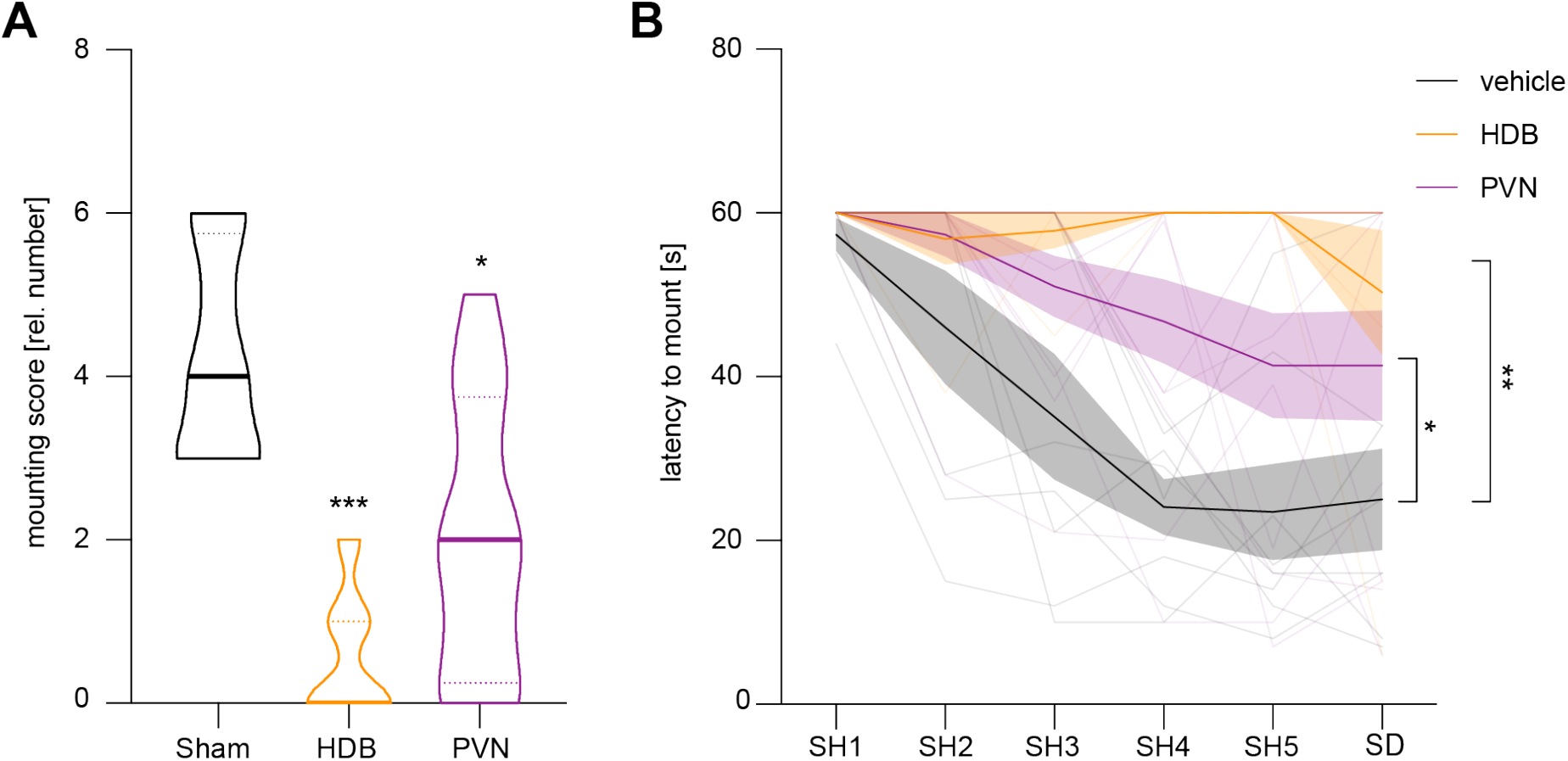
HDB- and PVN-silenced mice show impaired mounting behavior. **(A)** Bar diagram represents relative mounting score of vehicle-injected control (black, n = 8), HDB-silenced (yellow, n = 7) and PVN-silenced (purple, n = 12) mice, asterisks (***) indicate statistically significant differences in mounting score between groups calculated by 1-way ANOVA followed by Dunnett’s post-test. **(B)** Line diagram depicts the averaged latency (s±SEM) of test mice to mount a stimulus mouse. Asterisks (**) indicate statistically significant trial-independent differences in latency between groups calculated by 1-way ANOVA followed by Bonferonni’s post-test. *P < 0.05, **P < 0.01, ***P < 0.0001.

In sum, these results support the hypothesis of a sequential tri-synaptic basal forebrain circuitry linking social odor perception with non-social behaviors via HDB and placing the HDB as a central hub mediating both effects.

### Optogenetic silencing of HDB afferents within the OB prevents olfactory-mediated social behaviors

To further corroborate the important role of the HDB as a central hub guiding olfactory-mediated social behaviors, we again utilized miniSOG (Fig. 2 and 5) to selectively silence afferent projections from the HDB within the OB. Mice were injected with either rAAV expressing miniSOG (test group), or tdTomato (sham-injected control group) into the HDB, and light fibers were implanted into the GCL (Fig. 5A).

**Figure 5.**
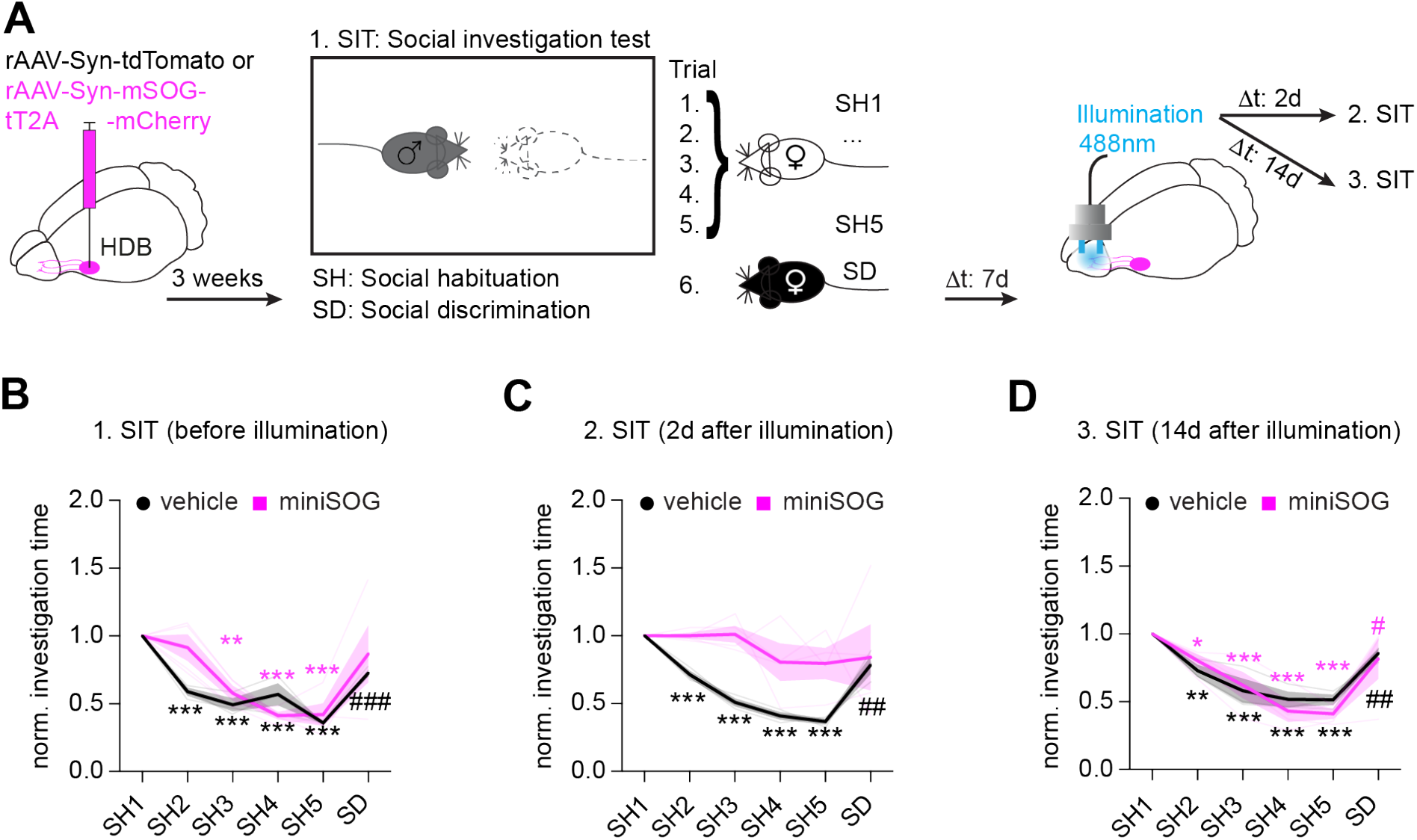
miniSOG-mediated silencing of HDB afferents within OB impairs olfactory-mediated social habituation/discrimination behavior. **(A)** Schematic of the experimental setup showing respective rAAV injections into HDB (left), SIT test (middle) and blue light illumination in the OB via chronically implanted light fibers (right). **(B-D)** Normalized investigation times (time the male spent investigating the female) of vehicle-injected control (black line) and miniSOG-injected mice (pink line) during SIT before blue light illumination (B), 2 days - (C) and 14 days (D) after the illumination. Repeated 1-way ANOVA followed by Dunnett’s posttest (SH1 to SH5, (*)) and t-test (SH5 to SD (#)). *p < 0.05, ***p < 0.01.

As already described before (see Fig. 2A), the SIT test was conducted before illumination (488nm) and subsequently two and 14 days after illumination (Fig. 5A). As expected, both vehicle-injected control and miniSOG test groups showed normal habituation and discrimination behaviors before illumination (Fig. 5B), excluding possible behavioral deficits due to chronic light fiber implantation. Notably, miniSOG-injected mice showed significant habituation and discrimination deficits two days after illumination. Fig. 5C clearly illustrates constant investigation times (pink line) during the entire test. Illumination had no effect on habituation and discrimination in the sham-treated control group (Fig. 5C, black line). Most importantly, the olfactory-mediated social behavioral deficits were completely reversed 14 days after illumination (Fig. 5D) indicating the reversibility of our approach.

Our results clearly show that selective silencing of HDB-OB projections leads to a complete loss of olfactory-mediated social habituation and discrimination behaviors. Together with our findings on odor habituation/discrimination deficits, these results place the HDB in a central position in modulating olfactory-dependent habituation/discrimination behaviors.

## Discussion

The OB is highly innervated by projections from the HDB. Chemogenetic and optogenetic silencing studies, as well as lesion studies suggested an involvement of the HDB in odor discrimination/habituation and odor-dependent social behaviors. These studies however used silencing strategies, which were not restricted to OB-innervating HDB projections only. Therefore, the simultaneous manipulation of the brain area’s targeted by the HDB like the hippocampus, piriform cortex and entorhinal cortex (Záborszky et al., 1986, Gaykema et al.; 1990) might have induced network effects that influenced the outcome of previous, especially investigations on the behavioral level. Thus, in the current study we performed specific miniSOG induced optogenetic silencing of HDB-projections within the OB. This leads to significant odor habituation/discrimination (OHOD) impairments as well as odor-dependent social discrimination deficits, thus ruling out a possible contribution of HDB-projections targeting other cortical and sub-cortical brain regions. Moreover, we used tissue clearing in combination with 3D confocal microscopy to reveal that HDB neurons innervate predominantly the GL and GCL in the OB. Additionally, we performed Ca^2+^-imaging of glomerular activity in awake head-fixed mice and showed that HDB silencing via virally expressed tetanus toxin light chain (TeTn) decreased odor-induced glomerular activity. The decreased glomerular odor responses correlated with impaired OHOD upon TeTn-mediated HDB silencing, corroborating previous results and indicating that HDB to OB projections have a modulatory effect on odor information processing in the OB. In addition, and since the HDB receives no direct input from the OB, we hypothesized that there might exist a poly synaptic loop connecting the OB with the HDB. To test this hypothesis, we used mono-transsynaptic retrograde tracing and found that HDB to OB projecting neurons receive monosynaptic input from the PVN of the hypothalamus. Accordingly, TeTn-mediated silencing of PVN led to odor-dependent social discrimination deficits that were comparable to HDB silencing effects. These results suggest the existence of a PVN-HDB-OB axis, that is critically involved in odor-dependent social habituation/discrimination tasks and may represent an anatomical feed forward link between the OB and the HDB.

### HDB silencing decreases glomerular activity which is associated with deficits in odor habituation and discrimination

The HDB is a basal forebrain region that extensively innervates the OB (Niedworok et al., 2012). By employing mono-transsynaptic retrograde tracing we demonstrate strong axonal input from the HDB predominantly targeting the granule cell and glomerular cell layers. Previous studies have shown that lesions of the HDB impair the animal’s ability to habituate to consecutive odor presentations (Paolini and McKenzie, 1993, 1996; Roman et al., 1993). This was confirmed by chemogenetic silencing of cholinergic and GABAergic HDB neurons, which was shown to disrupt discrimination of chemically similar odors (Nunez-Parra et al., 2013; Smith et al., 2015). Impaired odor discrimination possibly is a consequence of modulated odor-evoked OB output after activation of cholinergic and GABAergic HDB neurons (Bohm et al., 2020; Ma et al., 2012; Rothermel et al., 2014; Smith et al., 2015). Our TeTn-mediated silencing of HDB neurons resulted in decreased glomerular Ca^2+^-responses in awake mice during odor stimulation, underscoring the modulatory function of the HDB for odor-evoked responses in OB output neurons in awake mice. In line with our findings, previous studies using 2-photon Ca^2+^-imaging demonstrated the modulation of odor-evoked glomerular responses after electrical stimulation of the HDB (Bendahmane et al., 2016; Ogg et al., 2018). These studies investigating bulbar activity after HDB manipulation were conducted entirely in anesthetized mice. However, the utilization of anesthesia alters odor-evoked responses in neurons in the OB (Kato et al., 2012; Wachowiak et al., 2013). Thus, our study for the first time investigates odor-evoked glomerular responses after HDB manipulation in awake mice. HDB innervation to bulbar neurons seems to be essential in awake mice for accurate odor representation in the OB and consequently accurate odor discrimination. This is underscored by the impairment of odor discrimination and habituation behavior upon silencing of the HDB presented in this study. We used a behavioral habituation measuring the perceived similarity between odor stimuli thus allowing to investigate odor discrimination abilities without the formation of an association between odorant and reward (Cleland et al., 2002; Deiss and Baudoin, 1999; Guan et al., 1993). Consequently, the task is well suited for comparing behavioral results with imaging data that analyzes direct odor evoked neuronal activity (Cleland and Linster, 2002). Furthermore, preservation of cognitive abilities other than odor discrimination suggests that our test paradigm is not compromised by general cognitive deficits. These results are in line with previous findings showing impaired odor discrimination abilities after chemogenetic HDB manipulation (Nunez-Parra et al., 2013; Smith et al., 2015). Our observations also clearly indicate that HDB to OB projections significantly modulate Ca^2+^-responses within OB glomeruli *in vivo* which might lead to modulation of M/TC responses to odorants. The increase of odor-driven M/TC responses upon HDB activation was shown before in electrophysiological as well as Ca^2+^-imaging studies (Bendahmane et al., 2016; Bohm et al., 2020; Rothermel et al., 2014). Our observation of reduced odor-evoked glomerular Ca^2+^-responses upon HDB silencing supplements those studies and for the first time reveals glomerular responses after global silencing of the HDB in awake mice. The alteration of odor-evoked glomerular Ca^2+^-responses and impaired odor habituation/discrimination behavior after the same HDB manipulation supports our theory that HDB silencing affects the proper formation of bulbar odor representations interfering with olfactory habituation/discrimination behavior. However, the effect of global HDB-silencing and the resulting total loss of odor habituation/discrimination behavior was unexpected in its severity suggesting an instrumental role of HDB in odor processing besides its function as a global top-down modulator of olfactory perception.

### Optogenetic silencing of HDB afferents within the OB prevents odor habituation and discrimination behavior

Silencing the HDB by various means consistently resulted in odor habituation and discrimination deficits (Nunez-Parra et al., 2013; Paolini and McKenzie, 1993, 1996; Roman et al., 1993; Smith et al., 2015). Since the HDB not only innervates the OB, but also other cortical brain regions, including the hippocampus, piriform cortex and entorhinal cortex, network effects related to these brain regions cannot easily be excluded (Gaykema et al., 1990; Zaborszky et al., 1986). Therefore, a precise manipulation of exclusively HDB to OB projections would be necessary to exclude these network effects. However, up to now silencing specifically HDB to OB projections without affecting HDB-projections to other brain regions has not been reported. Here we utilized miniSOG to transiently silence specifically HDB projections within the OB’s granule cell layer upon blue light stimulation. This approach revealed a direct and reversible effect on odor habituation as well as odor discrimination behavior that was similar to that observed upon TeTn-mediated HDB silencing. These results clearly demonstrate that the HDB to OB projections are sufficient to mediate the deficits in odor habituation/discrimination behavior by changing OB activity patterns as we observed in glomerular *in vivo* Ca^2+^-response.

### The HDB is innervated by the paraventricular nucleus of hypothalamus and TeTn-mediated silencing of the PVN disrupts social habituation/discrimination behaviors

The fact that the HDB does not receive direct synaptic input from the OB suggests the existence of indirect, odor mediated activation pathways. To test the possibility of a polysynaptic loop connecting the OB with the HDB, we carried out retrograde mono-transsynaptic tracing experiments and found that neurons in the OB-projecting HDB neurons receive monosynaptic input from the PVN. Thus, we identified a novel PVN-HDB-OB axis that has a potential role in olfactory-mediated behaviors. The PVN also produces the neuropeptides oxytocin and vasopressin, which are relevant for social behaviors (Tang et al., 2020). Oxytocin positive neurons deriving from the PVN are shown to directly innervate the HDB in lactating rats (Knobloch et al., 2012). Furthermore, chemogenetic silencing of cholinergic HDB neurons impairs investigation of social odors (Smith et al., 2015). Thus, we hypothesized that this PVN-HDB-OB axis is involved in odor-dependent social behaviors. To test this hypothesis we virally expressed TeTn within the PVN and carried out a social habituation/discrimination test. Indeed, TeTn-mediated silencing of the PVN resulted in significant deficits in social habituation and discrimination. Notably and in analogy, TeTn-mediated silencing of the HDB showed similar effect on social behavior corroborating the notion that PVN and HDB form a functional axis. Along this line we could also show that TeTn-mediated silencing of either HDB or PVN led to major impairments in mounting behavior underlining the importance of this functional axis for successful reproduction. These results indicate that the PVN-HDB-OB axis is involved in different odor-dependent social behaviors. We conclude that the HDB acts as a relay station integrating incoming information with inputs from other brain areas to shape OB output depending on odor perception as well as the overall behavioral states. At this point it remains to be elucidated whether these functional pathways are mediated by separately acting, or overlapping circuits, or by distinctly organized structures within the HDB.

### Optogenetic silencing of HDB afferents within the OB disrupts social behavioral patterns

Our finding that TeTn-mediated silencing of the PVN and HDB affect social habituation and discrimination behaviors did not rule out the possibility of other network related effects due to PVN- or HDB-derived projections to other brain regions. To find out whether only HDB to OB projections are relevant for social habituation/discrimination behavior, we again used the miniSOG technology and selectively silenced HDB to OB projections. An impairment of social habituation and discrimination behavior was obvious two days after light stimulation and was recovered after two weeks. These results indicate that indeed specific projections from HDB to OB are required for normal social habituation and discrimination behavior. Moreover, these results reveal the importance of the HDB as a relay station to mediate odor-dependent social behaviors.Our approach of non-specifically silencing all HDB projections within the OB does not elucidate which neuronal types within HDB mediate the behaviorally relevant information. Since the chemogenetic silencing of cholinergic HDB neurons disrupt the identification of social odors (Smith et al., 2015), it is likely that they are relevant for odor-dependent social behavior. However, the involvement of other HDB populations in odor-dependent social behavior still needs to be investigated. In summary, we show that HDB projections innervating the OB are necessary for odor- and social habituation/discrimination behaviors. Furthermore, we identified the PVN as a new node within a PVN-HDB-OB axis relevant for social habituation and discrimination behaviors. The PVN-HDB-OB axis is part of a feedback loop transferring odor information from the OB via the HDB back to the OB with the HDB as a central nucleus involved in precisely modulating odor perception within the OB.

## Material & Methods

### Mice

For the experiments used mouse lines were C57BL/6J and the transgenic line *Thy1*-GCaMP6f GP5.5Dkim/J (Dana et al., 2014). All mice were housed separated by gender at a 12/12 h day/night cycle and received *ad libitum* access to food and water at all times (unless otherwise specified). All experiments were performed during the light phase and ahead of behavioral tests mice were allowed a 30 min acclimation phase to the testing room. Procedures were conducted in accordance with the National Institute of Health Guide for Care and Use of “Laboratory Animals” and animal welfare guidelines of Max-Planck Society and University Hospital Bonn, respectively, and were registered in the Regierungspräsidien Karlsruhe and Tübingen or approved by the North-Rhine-Westphalian government.

### Viruses

For the light induced silencing experiments (miniSOG) we used *rAAV-Syn-mSOG-t2A-mCherry, rAAV-Syn-tdTomato* (Addgene, Watertown, USA, plasmid number #50970). For mono-transsynaptic tracings we utilized, *RABVD-EGFP/EnvA, rAAV-CBA-RG* and *rAAV-CBA-tdTomato IRES TVA* exactly as described in (Niedworok et al., 2012). For tetanus toxin light chain mediated silencing we used *rAAV-Syn-TeTn-T2A-KO* (available upon request). To fluorescently label projections for tissue clearing and subsequent microscopic analysis we used *rAAV-Syn-EGFP* (Addgene, Watertown, USA, plasmid number #50465).

### Production, purification and characterization of rAAV vectors

A helper virus-free, three-plasmid-based strategy was applied to produce rAAV viruses (During et al., 2003). Co-transfection of AAV expression vectors and three AAV helper viruses (pRV, pH21, pFdelta6) was performed in HEK293 cells to generate viral particles of serotype 1/2. Viral particles were then purified using heparin columns. SDS-PAGE and infection of primary neurons were used to characterize new generated viruses.

Briefly, 48 – 72 hours after transfection fluorescent HEK293 cells were harvested in 9 ml PBS and spun for 10 min at 800 rpm and 4 °C. Each pellet was resuspended in 45 ml lysis buffer (20 mM Tris-HCl, 150 mM NaOH ph 8,0) and frozen at −20 °C over night. Upon thawing at RT, 6,6 µl Benconase (250 – 350 U/µl) and 2,25 ml 10 % NaDOC were added to each tube, in order to lysate HEK293 cells. Suspensions were incubated at 37 °C for 1 hour and spun for 15 min at 3000 x g and 4 °C. The supernatant was collected and frozen at −20 °C for heparin column purification. After thawing at RT, the suspension was centrifuged (15 min, 3000 x g, 4 °C) and the virus containing supernatant was loaded on a previously with lysis buffer pre-equilibrated 1 ml HiTrap Heparin HP column (SIGMA). Several washing steps later (increasing salt conditions, 100 – 300 mM NaCl), the virus was eluted under high-salt conditions (400 – 500 mM NaCl). Subsequently, the virus was concentrated and rebuffered with PBS in Amicon Ultra tubes and sterile filtered through a 0,2 µm Acrodisc column. Purification and integrity of viral capsid proteins (VP 1-3) were monitored on a coomassie-stained SDS/protein gel. The genomic titers were determined using the ABI 7700 real time PCR cycler (Life Technologies; Carlsbad, California, USA) with primers designed to bind the WPRE. Usually viral titers were 1 ×10^12^ /ml. rAAV-containing solution was aliquoted in 6 µl aliquots and stored at −70 °C.

### Production of modified, glycoprotein-deleted RABV for mono-transsynaptic tracing

BEK cells were plated at a density of ∼1.5 x 107. The following day, cells were transfected with 15 mg plasmid pCAGG/SAD-G by standard calcium phosphate transfection. 24 hours later, rabies virus SADΔG-mCherry or SADΔG-EGFP was added at a multiplicity of infection (MOI) of 3. 48 hours later the SADΔG-mCherry/EGFP containing supernatant was equally distributed onto four 15 cm plates containing pCAGGs/SAD-G (15mg/plate) transfected BHK cells (∼1.5 x 107 cells/plate). Two days later, the virus-containing supernatant was applied onto four 15 cm plates containing EnvARGCD cells (∼1.5 x 107 cells/plate) at an MOI of 1.5 for pseudotyping. 12 hours later, cells were trypsinized and replated onto eight 15 cm dishes. Pseudotyped rabies virus-containing supernatant was harvested 2 days later. The supernatant was spun at 2000 rpm at 4°C for 10 min. and subsequently filtered through a 0.45 mm filter (Nalgene; Thermo Fisher Sientific, Waltman, Massachusetts, USA). The filtered virus suspension was concentrated for 90 min. at 25000 rpm (SW28, 4°C) in an 80K ultracentrifuge (Beckman Coulter; Brea, California, USA). After centrifugation, the supernatant was discarded and the pellet was aspirated in ice-cold PBS (pH 7,4). Pseudotyped rabies virus-containing solution was aliquoted in 6 µl aliquots and frozen at −70°C. Virus titers were determined by serial dilution and overnight infection of primary cortical neurons that had been pre-infected with rAAV1/2 expressing TVA-IRES-EGFP/Cherry under the control of human synapsin promotor / enhancer. Three days later, the number of red/green fluorescent, SADΔG-mCharry/EGFP containing neurons were counted. Biological titers on TVA-expressing neurons for SADΔG-mCherry/EFFP for in vivo injections were ∼2.5 x 10^7^/ ml.

### Stereotaxic injections

Mice were deeply anaesthetized intraperitonially using a mixture of ketamine hydrochlorid (90mg/kg) and xylazine (5mg/kg) or ketamine (0.13 mg/g) and xylazin (0.01 mg/g) in PBS pH, 7.4. For window implantation (s. below) an analgesic (0.05 mg/kg burprenorphin) and anti-inflammatory drug (0.2 mg/kg bodyweight dexamethasone) were delivered by s.c. injection. Eyes were protected with eye ointment (Bepanthen, Leverkusen, Germany) and body heat was maintained using a heating pad (Stoelting’s Rodent Warmer X2, Stoelting Europe, Dublin, Ireland). After fixation in a stereotaxic frame (Narishige, Tokyo, Japan or David Kopf Institute, Tujunga, California, USA) skin was shaved, disinfected with 70 % ethanol and numbed with 1 % lidocaine hydrochloride. The skull was exposed with a vertical incision in the skin and the remaining tissue was removed. Levels of the anterior-posterior and medio-lateral plain were manually adjusted by comparing bregma to lambda, and left to right. Injection sites were determined from bregma (tab. 3.1) and small craniotomies performed accordingly. A 33 or 34 gauge beveled needle fitted to a 10 µl syringe containing the viral solution was inserted at target depths (tab 3.1) relative to brain surface and virus-containing solution was injected either uni- or bilaterally at a speed of 100 nl/min by a microprocessor-controlled minipump (World Precision Instruments, Sarasota, Florida, USA). The virus was allowed to diffuse into the tissue for 10 min. before slowly removing the injection needle. Subsequently the skin was sutured (PERMA-HAND Silk Suture 3-0) and maintaining body temperature was ensured until recovery. Treatment with analgesics (buprenorphine (0.05 mg/kg) or and ketoprofen (5 mg/kg)) was conducted for 3 days post-surgery. Synaptic transmission from HDB was silenced by injecting a total of 500 nl of *AAV1/2-hSyn-TeTn-2A-KO* (available upon request). For miniSOG-mediated, light induced silencing 1 µl of *rAAV-Syn-mSOG-t2A-mCherry*, or *rAAV-Syn-tdTomato* (Addgene, Watertown, USA, plasmid number #50970) was per OB was injected.

### Light fiber implantation

For the optogenetic experiments (miniSOG-mediated silencing), the light fibers (Dual Fiber-optic Cannula DFC_400/430-0.37_1.5mm_GS1.7_B45, Doric Lenses Inc, Canada) were implanted over the center of the OB of adult male C57Bl/6 mice (Charles River) ∼one week after virus injection. Briefly, after exposing the skull, the area located over OB was covered with light curable dental adhesive (OptiBond FL, Kerr, USA) and small craniotomies were performed bilaterally over OB. The light fiber was lowered in OB under the control of a custom-made holder and fixed with blue light curable dental cement. Subsequently the skin was sutured (PERMA-HAND Silk Suture 3-0). The body temperature of the mouse was controlled until recovery. Mice were treated with analgesics (0.05 mg/kg buprenorphine) for 3 consecutive days after the surgery and were allowed to recover at least 3 weeks before the experiment.

### Optogenetic silencing of HDB-OB projections

For optogenetic silencing of HDB-OB projections, the mice were lightly anesthetized with a mixture of Fentanyl, Midazolam, and Medetomidin injected intraperitoneally (0.02/2.0/0.2 mg/kg). The fiber cannulas were connected via dual fiber-optic patch cord (DFP_400/440/900-0.22_3m_GS1.7_2FC, Doric Lenses Inc, Canada) to a 488 nm laser. The OB were illuminated 5 times for 5 min with 1 min ITI with an effective laser power of 30 mW at the end of the glass fibers. Behavioral experiments were conducted 2 and 14 days after illumination.

### Window implantation

For *in vivo* imaging in the olfactory bulb a chronic cranial window was implanted above the OB approximately one week after virus injection. Anesthesia was induced by i.p. injection of ketamine/xylazin (0.13/0.01 mg/g bodyweight). Mice were injected with burprenorphin (0.05 mg/kg bodyweight) and dexamethasone (0.2 mg/kg bodyweight). Mice were fixed in a stereotaxic frame and body temperature was controlled using a heating pad (Stoelting’s Rodent Warmer X2, Stoelting Europe, Dublin, Ireland). Eye ointment (Bepanthen, Leverkusen, Germany) protected the eyes throughout the whole surgery. After removing the skin, the skull was covered with a light curable dental adhesive (OptiBond FL, Kerr, USA). Subsequently, a craniotomy was carried out above the olfactory bulb using a dental drill (2×3 mm). The bone and the *dura mater* were carefully removed. The craniotomy was sealed by a coverslip fixed with UV curable glue (GRADIA^®^ DIRECT Flo, GC EUROPE N.V., Leuven, Belgium). A metal bar (Luigs and Neumann, Ratingen, Germany) for head fixation was attached posterior to the cranial window. Mice were treated with analgesics (0.05 mg/kg burprenorphin) for 3 consecutive days after surgery and allowed to recover at least 3 weeks before imaging.

### Behavioral Analysis

For behavioral analysis adult male C57Bl/6 mice from Charles River laboratories were used. All ‘test’ mice were bilaterally injected with a rAAVs expressing the respective transgenes. ‘Control’ mice were vehicle-injected with a rAAV expressing EGFP.

### Puzzle box

The puzzle box test was used to study general abilities and operant learning in TeTn-injected and vehicle-injected control mice. The apparatus consisted of a Plexiglas box, divided by a black-painted Plexiglas into two compartments. The small black-painted goal compartment contained home-cage bedding in a 10 cm petri dish, and covered with a lid. The open start compartment was white painted and brightly illuminated (∼ 1,000 lux) by a neon lamp. The entrance to the goal compartment was blocked by different kinds of barriers (see below). In order to reduce environmental cues, which could bear additional anxiety, the apparatus was placed inside a Faraday cage covered by black drapery.

A trail started by placing a mouse into the start compartment with its head facing the opposite site of the entrance. Trial duration was 3 min for the first seven trials and 4 min for the last three trials. Trials were separated by either a short (1 min.) or a long (24 hours) inter-trial interval (ITI). At the end of each successful trial mice were allowed to stay in the goal compartment for approximately 1 min and then returned to their home cage. The goal compartment was cleaned with water after a mouse completed its daily trials, while the start compartment was cleaned between each trial.

*Following barriers were used to study operant learning and general abilities in mice:*

1. door barrier (trail 1): mice had to enter the goal compartment simply by running passing an entrance within 3 min
2. underpass barrier (trail 2 - 4): to enter the goal compartment mice had to go through an opened underpass within 3 min
3. sawdust barrier (trail 5 - 7): the underpass was filled with sawdust requiring mice to burrow through in order to enter the target compartment within 3 min
4. plug barrier (trail 8 - 10): the underpass was plugged by a cardboard. In order to enter the goal compartment mice were required to remove the plug within 4 min

### Buried food test (BFT)

Ability to smell volatile odors was studied in the buried food test as described in (Yang and Crawley, 2009). It relies on the natural tendency of mice to use olfactory cues for foraging. Overnight-fasted mice were placed in a clean standard plastic cage containing 3 cm deep fresh cage bedding and allowed to explore the new environment for two minutes. Then they were transferred to an empty clean cage for 1 minute. During this time the food stimulus (piece of oat cake), which mice were habituated to on previous days, was buried in the sawdust in random corners of the bedding-containing cage. Upon re-introduction to the cage mice had to find the buried food within 15 minutes. Latency to locate and consume the food was measured.

### Olfactory habituation /discrimination test (OHOD)

In order to assess the ability of injected animals to detect and distinguish similar and different odors, mice were tested in a simple olfactory habituation/discrimination paradigm. It relies on the animals’ tendency to investigate novel scents. By repeated presentation of the same odor stimulus a progressive increase in latency and decrease in olfactory investigation indicates olfactory habituation. Dishabituation is defined by a reinstatement of latency and investigation when a novel odor is presented. The test was performed in the open field arena containing a 10 cm petri dish equipped with bottom-placed Whatman filter paper (25 mm diameter, Sigma-Aldrich, Steinheim, Germany) and filled with fresh bedding. It consisted of two habituation trials followed by sequential presentations of 2-phenylanaline (3 times) and vanillin (1 time) soaked on the Whatman filter paper. 100 µl of a 1:1000 dilution of 2-phenylethanol was utilized for experiments. On the last stimulus trial (dishabituation) vanillin was applied to the filter paper. Trail duration was either 120, or 240 sec. and inter-trail interval (ITI) was 2min. Latency to enter the petri dish and duration of investigation was automatically measured using NOLDUS software.

### Social habituation/discrimination test (SIT)

Social interaction behavior was studied in a simple social habituation/discrimination paradigm. It relies on the ability of mice to recognize familiar conspecifics. Familiarization (habituation) is indicated by a decline in investigation measured in repeated encounters between the same mice. While discrimination (dishabituation) is indicated by re-emergence of vigorous investigation when a new stimulus mouse is presented. The Social habituation/discrimination test was performed according to (Winslow, 2003). Briefly, each test and control (sham-injected, rAAV-Syn-tdTomato; 500nl/hemisphere) mouse encountered on five repeated 1 min trials the same stimulus mouse in its home-cage. On the sixth “discrimination” trial a new stimulus mouse was presented for one minute. Each trial was separated by a 10 min. ITI. As stimulus mice 10-weeks old ovariectomized (OVX) C57/Bl6 female mice were used. Ovariectomy was performed 4 weeks before testing. Encounters were video recorded and analyzed for investigation duration and social interactions.

### *In vivo* 2P Ca^2+^-imaging

2-Photon awake Ca^2+-^imaging was conducted head-fixed on a linear track. Prior to imaging mice were habituated to head fixation and setup for 3 sessions á 1 h. For the imaging session mice were placed on the linear treadmill and head-fixed to prevent movement of the brain. An olfactometer (Two Channel Dilution Olfactometer, MED Associates Inc., Fairfax, USA) was placed directly in front of the nose. A video camera was used to record the mouse behavior during the experiment (Sumikon PX-8262-919). The microscope objective (N16XLWD-PF, 0.8 N/A, Nikon, Düsseldorf, Germany) was positioned above the cranial window and changes in Ca^2+^-transients recorded using a galvo-resonant scanner (Thorlabs, Newton, USA) on a two-photon microscope (ThorLabs, Newton, USA) provided with a Ti:Sa laser (Coherent, Santa Clara, USA) tuned to 920 nm. Each trial consisted of a 2 min Ca^2+^-imaging with an odor-stimulation after 50 s lasting for 5 s. The frame trigger of the microscope was recorded with an ITC-18 board (HEKA, Ludwigshafen/Rhein, Germany) and used to synchronize the imaging with the odor stimulation (IGOR Pro software, WaveMetrics, Portland, USA). Presented odors were rose (undiluted 99% 2-Phenylethanol, Sigma-Aldrich, Darmstadt, Germany) and vanillin (59.15 mM in 0.5 % EtOH in H_2_O, Sigma-Aldrich, Darmstadt, Germany).

### Data analysis for *In vivo* Ca^2+^-imaging

Acquired images were registered in x,y-dimension using the Lucas-Kanade algorithm (1981). ROIs for analysis of glomerular Ca^2+^-transients were drawn manually. The mean gray values in selected ROIs were extracted using FIJI. Further analysis was conducted using a MATLAB® (Mathworks, Natick, USA) pipeline. In brief, the pipeline applied a low-pass Butterworth filter (cutoff 1 Hz), calculated ΔF/F by subtracting the baseline fluorescence F_0_ and dividing by F_0_ and fitted a binominal curve to the data points recorded 10 s after stimulus onset. F_0_ was defined as mean of the 10th percentile of values from 10 – 45 s of each time-lapse. Following the curve-fit transients were detrended and trials averaged. From those averages area under the curve was calculated as integral of the calculated binominal curve.

### Microscopy

Mosaic, multiple-channel fluorescent overview images were generated at the LSM5 (Zeiss) using a 5x or a 10x magnification objective. A built-in algorithm of the Zeiss AxioVision control software stitched single images to assemble mosaic image. Confocal images were obtained using a customized Leica SP5 confocal microscope (Leica Microsystems) using a 10x or 20x magnification objective to evaluate co-labeling of rabies virus positive and immunochistochemically stained fluorescent cells in HDB and PVN. A built-in algorithm of the Leica control software stitched single images to assemble mosaic-merge images (Version 2.6.0.7266, Leica Microsystems). All pictures are maximal projections of 70 μm thick tissue slices.

### Tissue clearing

For clearing of whole adult mouse brains, specimens were treated as previously described. Briefly, tissue was dehydrated using triethylamine pH 9.5 adjusted tert-butanol-based ascending alcohol series at 30 °C and mounted in a triethylamine pH 9.5 adjusted benzyl alcohol / benzyl benzoate mixture (BABB) (Schwarz et al., 2015).

### 3D reconstruction of imaged data

For generation of traverse Z-projections, the maximum intensity projection algorithm of Fiji was used (http://pacific.mpi-cbg.de/wiki/index.php/Fiji). The representation of the generated data as a 3D object was performed using the Surpass view in Imaris while virtual sections of varying thickness were generated using the extended feature of the slice tool (Bitplane). Data processing was performed on a custom-built power workstation holding two Xeon E5-2667v3 CPUs, 512 GB memory and a NVidia Titan-X GPU.

### Statistics

Statistical analysis was carried out using GraphPad Prism 5 or 7 (GraphPad Software Inc., La Jolla, USA). Data was tested for normality using the Shapiro-Wilk normality test and glomerular activity (Sham- vs TeTn-treated) was compared applying the Mann-Whitney test with uncorrected alpha (desired significance level) set to 0.05 (two-tailed) (* < 0.05, ** < 0.01, *** < 0.001, ****p < 0.0001,). For behavior test differences within groups were determined either by repeated measures ANOVA or by Friedman Test (as a non-parametric test with repeated measures). When ANOVA revealed a statically significant difference pair wise multiple comparison procedures were used for post hoc testing (Bonferroni’s, Dunnett’s or Dunn’s). To determine differences between groups either the unpaired t-test (normal distributed, equal variance) or two-way ANOVA was used depending on number of groups. Regardless of the type of test chosen, uncorrected alpha (desired significance level) was set to 0.05 (two-tailed). Data are displayed as the mean and standard error of the mean (SEM) or 95 % percentiles.

## Data Availability

The datasets generated and analyzed during the current study are available from the corresponding authors on request.

## Code Availability

The code used for analyzing the data in the current study is available from the corresponding authors on request.

## Acknowledgements

This work was supported by grants from the Deutsche Forschungsgemeinschaft SFB 1089 C01 MF and B06 MF/MKS as well as SPP2041 to MKS. Further, this work was supported by the DZNE, the ERA-NET MicroSynDep and MicroSchiz projects, as well as by the European Union Horizon 2020 ERC CoG MicrSynCom to MF. We would like to thank Heinz Beck for continuously supporting the progress of the project with scientific discussions and suggestions. We would like to thank Karl-Klaus Conzelmann for support and supply with modified rabies virus. We would also like to thank Jonas Doerr for help with confocal 3D imaging and IMARIS software processing and Lydia Fischer for excellent technical support. We thank P. Thevenaz and Erik Meijering for the development of the ImageJ plugins “stackreg” and “TurboReg”. We thank the Light Microscope and Animal Facilities of UKB and DZNE for constant support.

## Author contributions

Conceptualization, M.K.S and M.F.; Methodology, M.M., and I.S.; Formal Analysis, I.S., M.M., M.Mi., F.M. and J.W.; Investigation, I.S., M.M.; Resources, J.S. F.M. and M.Mi.; Writing – Original Draft, M.K.S, M.F., M.M., I.P. and I.S.; Writing – Review & Editing, M.K.S., M.F., I.S., M.M.; Visualization, M.M., I.S; Supervision, M.K.S. and M.F.; Funding Acquisition, M.K.S. and M.F.

## Supplemental Information

**Supplementary figure 1.**
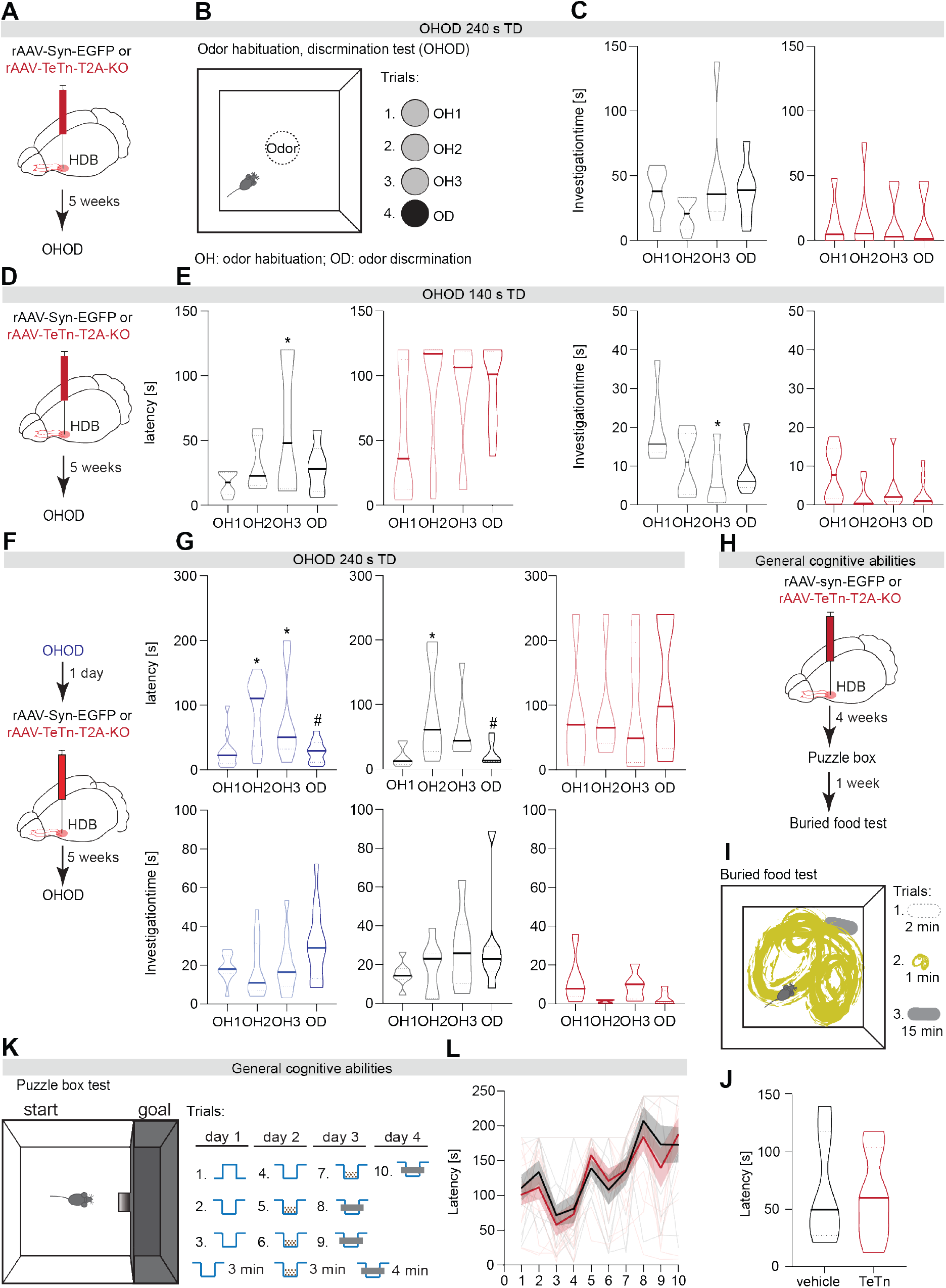
Tetanus toxin-mediated silencing of HDB disrupts odor habituation and discrimination without affecting general cognitive abilities. **(A)** Schematic of the stereotaxic injection. **(B)** The odor habituation/discrimination test (OHOD) performed in an open-field arena (60 x 60 cm). The circle in the middle of the arena indicates a petri dish containing a paper with odorant covered with fresh bedding. 2-phenylethanol (rose odor) and vanillin were used in habituation and discrimination trials, accordingly. **(C)** Investigation time (time spent at petri dish, 120 or 240 sec trial duration and 2 min. TD) of control and HDB silenced mice during odor habituation and discrimination trials (see also Fig. 1). **(D)** Schematic of the stereotaxic injection. **(E)** Latency (time to reach the petri dish) and investigation time of control and HDB silenced mice during odor habituation and discrimination trials with shorter trial durations. **(F)** Schematic stereotaxic injection and sequence of experiments. **(G)** Latency and Investigation time of control and HDB silenced mice during odor habituation and discrimination trials before (blue) and after TeTn virus injection. **(H)** Schematic of test sequence and stereotaxic injection. **(I)** Buried food test was conducted in home cage. The ellipse indicates a hidden food pellet. **(J)** Latency in finding the food pellet of control and HDB silenced mice during buried food test. **(K)** The puzzle box test paradigm. Left: the arena is divided by black-painted Plexiglas into the start and goal compartments. Right: a timeline of the experiment; the compartments are connected by a door opening (trial 1), open underpass (trials 2-4), underpass blocked by sawdust (trials 5-7), or underpass blocked by a cardboard plug (trials 8-10). **(L)** Latency (time to enter the goal compartment) of control and HDB silenced mice. 1-way ANOVA followed by Dunnett’s posttest (*) and t-test (#) of sham-injected (n=6) and TeTn-injected (n=6) mice. #, *p < 0.05, **p < 0.01. All data represented as mean ±SEM (transients) or median ±quartiles (violin plots). HDB: horizontal limb of the diagonal band of Broca; MOB: main olfactory bulb; GL: glomerular layer; EPL: external plexiform layer; GCL: granule cell layer.

**Supplementary Figure 2.**
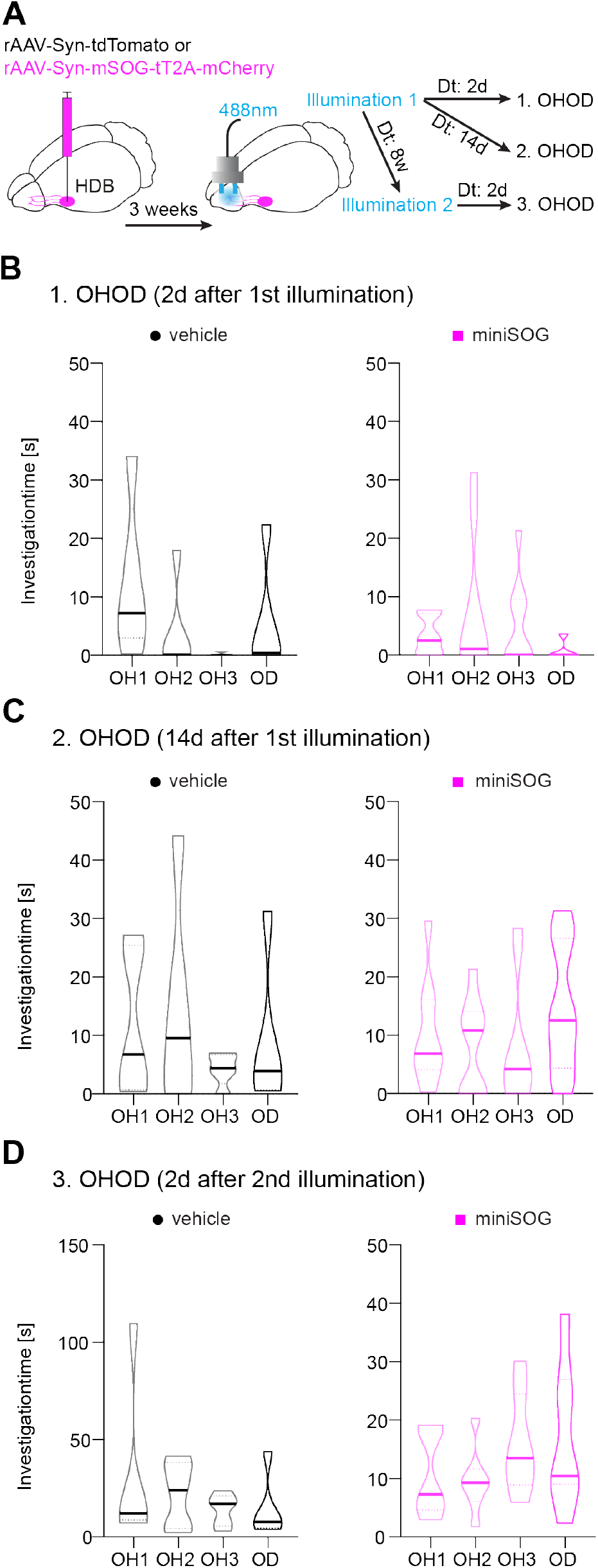
miniSOG-mediated, light-induced silencing of HDB afferents within OB impairs odor habituation/discrimination behavior with respect to investigation time. **(A)** Timeline of the experiment. The odor habituation/discrimination task (OHOD) was performed as described in Fig 1N, O. **(B-D)** Investigation time of vehicle-injected control and miniSOG-silenced mice during odor habituation and discrimination trials 2 days (B) and 14 days (C) after the first illumination, as well as 2 days (D) after the second illumination. Repeated 1-way ANOVA followed by Dunnett’s posttest (SH1 to SH3, (*)) and t-test (SH3 to SD (#)). *p < 0.05, **p < 0.01.

